# Improved maximum growth rate prediction from microbial genomes by integrating phylogenetic information

**DOI:** 10.1101/2024.10.03.616540

**Authors:** Liang Xu, Emily Zakem, JL Weissman

**Affiliations:** Department of Global Ecology, Carnegie Institution for Science, Stanford, CA, USA; Institute for Advanced Computational Science, Stony Brook University, Stony Brook, NY, USA; Department of Ecology and Evolution, Stony Brook University, Stony Brook, NY, USA; Department of Biology, The City College of New York,New York, NY, USA

## Abstract

Microbial maximum growth rates vary widely across species and are key parameters for ecosystem modeling. Measuring these rates is challenging, but genomic features like codon usage statistics provide useful signals for predicting growth rates for as-yet uncultivated organisms, though current predictors often show high variance. To improve accuracy, we integrate phylogenetic signals, leveraging the evolutionary relationships among species to refine trait predictions. We present *Phydon*, which combines codon statistics and phylogenetic information to enhance the precision of maximum growth rate estimates, especially when a close relative with a known growth rate is available. We construct the largest and most taxonomically broad database of temperature-corrected growth rate estimates for 111,349 microbial species. The results reveal a bimodal distribution of maximum growth rates, resolving distinct groups of fast and slow growers. Our hybrid approach advances the accuracy of genome-based growth rate predictions and presents a new framework for accurate genome-based trait prediction.

## Introduction

Microbes are crucial players in global nutrient cycles, and their maximum growth rates are key parameters in ecosystem models [1-3]. Traditionally, maximum growth rates are inferred from measurements using laboratory isolates or field-based methods, such as nutrient uptake experiments [4, 5] and peak-to-trough measurements [6, 7]. However, accurately measuring rates for many species poses significant challenges in both laboratory and field settings [8]. Only a small fraction (less than 1%) of bacterial and archaeal species from any given environment has been successfully cultured [9, 10]. Even among these cultured species, maximum growth rates vary widely, with population doubling times ranging from minutes to days across species and culture conditions [9, 11-13], adding further complexity to measurement and cultivation efforts.

As a powerful alternative to cultivation for hard-to-grow species, genomic information can be leveraged to estimate the maximum growth rate of an organism. Maximum growth rate—how fast a population can grow under optimal conditions—is a broad indicator of an organism’s overall evolutionary strategy and a key parameter for describing population dynamics. Several genomic features have been linked to maximum growth rates, including rRNA operon copy number [14-17], tRNA multiplicity [18, 19], replication-associated gene dosage [20, 21] and codon usage biases (CUB) [18, 22, 23]. Among these, CUB has shown the strongest correlation with growth rates [11, 12]. In fast-growing species, highly expressed genes tend to preferentially use certain synonymous codons, a bias that arises from the need for efficient translation. This optimization ensures the rapid production of proteins [22]. The robustness of CUB has been demonstrated even when extrapolating predictions across different phyla [11].

Although codon usage bias (CUB) is a widely used genomic predictor of maximum growth rates, the resulting estimates can exhibit considerable variance [11]. This variability may stem in part from the fact that traits such as growth are influenced by multiple genetic factors while CUB captures only one aspect of this complexity. Therefore, while CUB reflects evolutionary optimization for rapid translation and, by extension, rapid growth, its precision in estimating growth rates is theoretically limited. To improve accuracy, additional signals can be incorporated. One such signal is phylogenetic relatedness: closely related species tend to exhibit similar trait values due to their shared evolutionary history [24] and vertical gene inheritance [25]. This phenomenon, known as phylogenetic signal [26, 27], offers a complementary approach to CUB and can help reduce the variance in CUB-based growth rate predictions.

The simplest model for trait inference from phylogenetic trees estimates a species’ trait based on the trait of its nearest neighbor in the phylogenetic tree, leveraging the tendency for phylogenetically related organisms to exhibit similar phenotypes [28]. More advanced approaches involve specifying models of trait evolution, commonly using a Brownian motion framework, where traits values can be estimated for a query species based on its position in the tree and distance to neighboring species [24, 28-35].

More sophisticated trait evolution models, beyond the Brownian motion model, exist [32-34, 36], but they often require additional information, such as convergent trait values or eco-evolutionary timescales. Of course, the accuracy of all phylogenetic prediction methods generally decreases as phylogenetic distance increases, depending on how strongly conserved the trait is across evolutionary time.

Microbial traits vary widely in their degree of phylogenetic conservation. Martiny et al. [27] developed a phylogenetic metric to quantify this conservatism and estimate the clade depth at which organisms share a given trait. They found that over 90% of functional traits in their data were significantly non-randomly distributed. However, the clades sharing these traits were generally shallow, suggesting a moderate degree of phylogenetic conservatism. As a result, the accuracy of utilizing phylogenetic structure to estimate trait values remains uncertain and may vary by trait and context [37]. Typically, complex traits involving multiple genes (e.g., photosynthesis or methanogenesis) tend to exhibit stronger phylogenetic conservatism than simpler traits, such as the consumption of a specific carbon source [27, 37]. This raises the question of whether phylogenetic relationship can reliably predict maximum growth rates, which are determined not by any one gene or set of genes, but rather as a complex outcome of a variety of genomic factors. Walkup et al. [26] assessed phylogenetic-based predictions of bacterial growth rates across environments and found that phylogenetic relationships accounted for only 38% variation in maximum growth rates across ecosystems. Moreover, such tools will only work well when a high-quality reference database of species with known trait annotations already exists for a given environment. Thus, the effectiveness of phylogenetic prediction methods depends heavily on the quality of trait data and phylogeny, as well as the strength of the phylogenetic signal.

In this study, we aim to enhance the accuracy of estimating maximum growth rates by integrating codon usage bias (CUB) with phylogenetic relatedness to create a hybrid approach for trait prediction. We evaluated the performance of a genomic CUB-based method (gRodon; [11]) against two phylogenetic prediction models: the nearest-neighbor model (NNM) and the phylogenetic independent contrast-based Brownian motion model (Phylopred). To ensure robust evaluation, we tested model performance across a range of phylogenetic distances. Our analysis identified a threshold in both phylogenetic distance and growth rates where the predictive power of CUB and phylogenetic signals intersect. Specifically, the CUB-based approach outperformed phylogenetic methods for distantly related query genomes, while phylogenetic models excelled for closely related sequences. Based on these findings, we developed a novel R package, Phydon, which synergistically combines both approaches. Our results demonstrate that Phydon significantly improves the accuracy of maximum growth rate estimations for microbial genomes, particularly for faster-growing organisms and when a close relative with a measured maximum growth rate is available.

## Results & Discussion

### A Phylogenetically Informed Model for Maximum Growth Rate Prediction

The maximum growth rates of the species in our training dataset exhibit a moderate phylogenetic signal, as indicated by a Blomberg’s *K* statistic [28] of 0.137 and a Pagel’s λ statistic [38] of 0.106 with p-value < 0.0072 for bacteria species, and a Blomberg’s *K* statistic of 0.0817 and a Pagel’s λ statistic of 0.17 with p-value < 0.0055 for archaea species. For reference, a value approaching or exceeding 1 indicate strong phylogenetic conservatism, while a value of 0 indicates no phylogenetic signal. These values suggest that while there is some degree of phylogenetic conservatism (Fig. 1), it is not overly strong. This makes the dataset well-suited for developing a method that balances genomic and phylogenetic factors. We further explored how different prediction methods perform under varying conditions to identify the most effective approach for improving growth rate predictions. The phylogenetic distance between the training and test datasets is a critical factor in evaluating the performance of the two methods. To assess this, we successively divided the phylogenetic tree into two groups (training and test) based on varying phylogenetic distances, which is known as phylogenetic blocked cross-validation analysis [39] (Fig. 2). Each method was trained on each training dataset, and its performance was evaluated using each test dataset respectively. In doing so, the ability of each model to extrapolate to new taxonomic groups not in the training data was tested directly. For further details on the analysis design, we refer readers to the Methods section.

**Figure 1.**
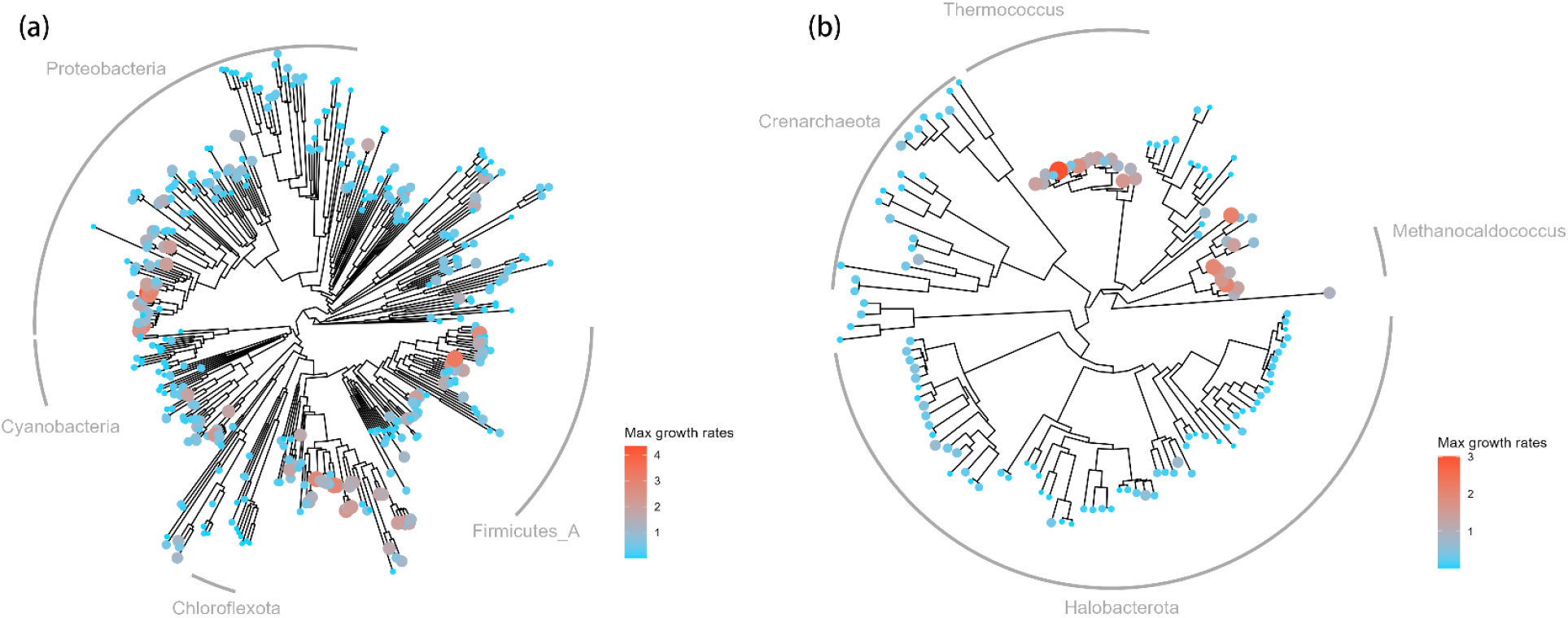
Phylogenetically conserved patterns in the maximum growth rates of bacteria (a) and archaea (b). Colors indicate the magnitude of maximum growth rates, with faster-growing species represented by warmer tones. Among bacteria, certain groups within *Proteobacteria* and *Firmicutes_A* exhibit higher growth rates, while in archaea, the genera *Thermococcus* and *Methanocaldococcus* are notable for their rapid growth.

**Figure 2.**
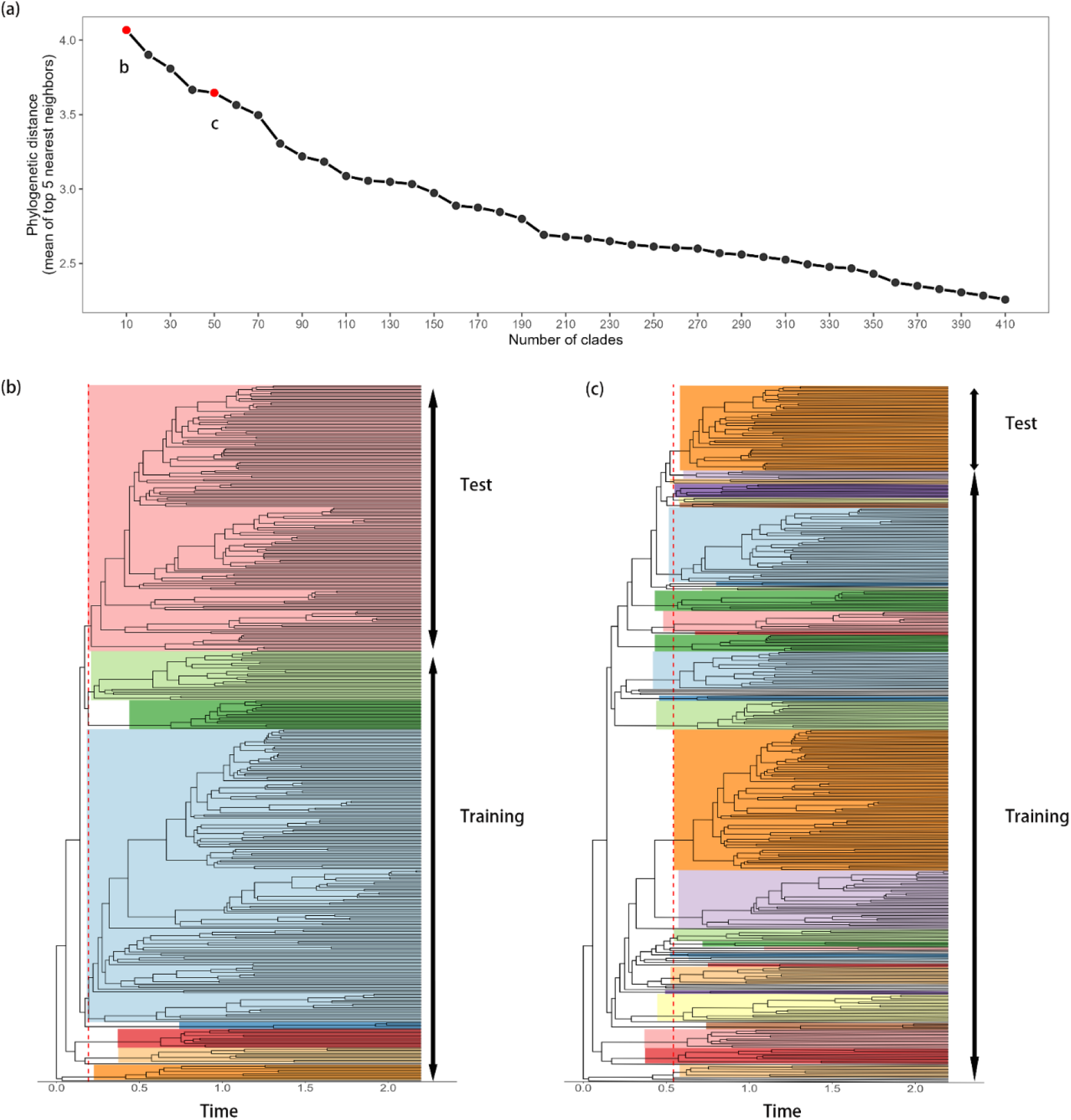
a) The phylogenetic distance between the training and test data sets when cutting the tree at different heights; b), c) The illustration of the phylogenetic trees cut at two time points where the tree is separated into 10 versus 50 clades for blocked cross validation.

The gRodon model consistently provides accurate directional predictions of bacterial growth rates across the tree of life, as demonstrated by a stable mean squared error across varying phylogenetic distances (Fig. 3a). This finding supports the notion that codon usage bias serves as an effective genomic proxy for bacterial growth rates [11, 12], capturing selective pressures on genomes over evolutionary time [18, 19, 22, 40]. Additionally, our phylogenetic blocked cross-validation analysis (see Methods, Fig. 2) indicates that the relationship between CUB and the maximum growth rates generalizes well across different clades. However, we also observed significant variance in the growth rate estimates, which persists even with decreasing phylogenetic distance between training and test sets (Fig. S1). This suggests that while CUB is a valuable predictor, additional factors beyond codon bias influence bacterial growth rates.

**Figure 3.**
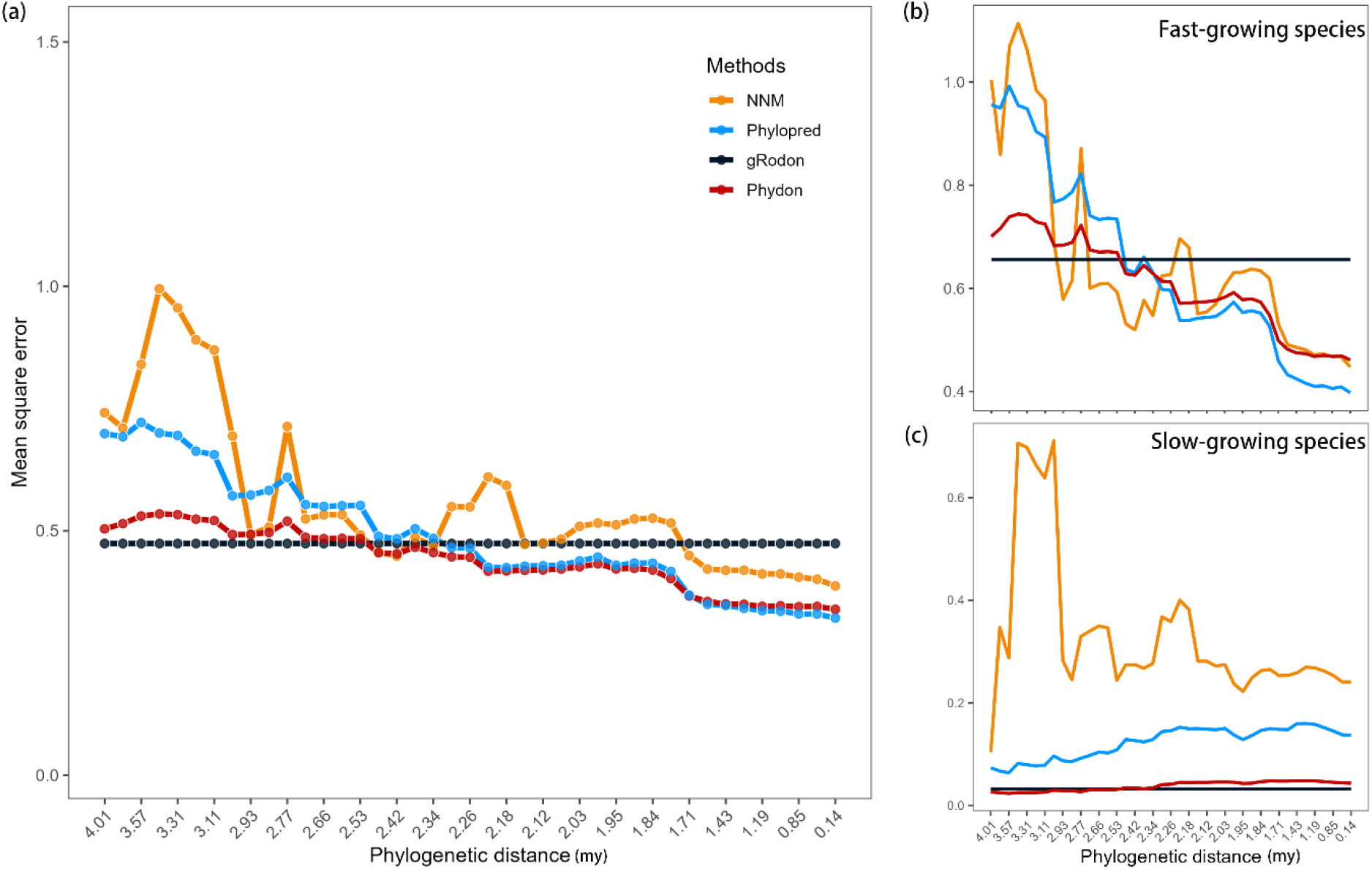
a) The prediction error of the four methods varies with phylogenetic distance; (a) Blocked cross-validated error estimates for each method when folds for cross-validation were determined phylogenetically with the cutoff of phylogenetic distance varied b-c) the MSE scores for the slow-growing species (the doubling time > 5 hours) and fast-growing species (the doubling time < 5 hours).

Phylogenetic prediction methods show increased accuracy as the minimum phylogenetic distance between the training and test sets decreases. As shown in Figure 3a, the mean squared error (MSE) for both the NNM and Phylopred models decreases significantly as the minimum phylogenetic distance between the training and testing data narrows from 4.01 *my* to 0.14 *my*. We identified specific phylogenetic distance thresholds below which the MSE of these phylogenetic models falls below that of the gRodon model. For Phylopred, this threshold is 2.42 *my*, while for NNM, the MSE initially drops below gRodon’s at 2.53 *my*, but then fluctuates around gRodon’s performance until the phylogenetic distance drops below 1.73 *my*. These results suggest that Phylopred and NNM have distinct thresholds, with Phylopred showing more stable and superior performance. Based on this, we chose the Phylopred model to develop a combined approach with the gRodon model.

Interestingly, we observed divergent performance patterns between the gRodon and Phylopred models for fast-growing and slow-growing species (Fig. 3bc). For slow-growing species, the gRodon model consistently outperforms the Phylopred model across all phylogenetic distances (Fig. 3c). In contrast, the Phylopred model shows superior performance over the gRodon model for fast-growing species as the phylogenetic distance decreases (Fig. 3b).

We sought to build a predictive model that combined the strengths of these two approaches, taking into account which model would most likely work best for a given organism. To achieve this, we developed a weighted predictor where the weight of each model was determined by both the distance of the query genome to the model training set and by the gRodon estimate of growth rate (as a rough estimate of whether an organism was likely to be a fast or slow grower, see Methods). This ensemble model, implemented in the R package Phydon, demonstrates superior predictive accuracy compared to the individual models on average (Fig.3). Specifically, the Phydon model achieves lower Mean Squared Error (MSE) scores under most of the phylogenetic distances, while similar MSE scores compared with that of the gRodon model were observed when phylogenetic distances are large. Additionally, the variance of the Phydon’s predictions is lower than that of alternative approaches (Fig. S1).

We defined the weight parameter *P* as a continuous value between 0 and 1 (see Method), ensuring that Phydon estimates always incorporate information from both phylogenetic relationships and genomic statistics. Alternatively, if the weight parameter *P* is treated as a binary variable, the model consistently achieves the lowest MSE scores across all test scenarios. However, this approach leads the model to discard information from one source entirely, favoring either phylogenetic relationships or genomic statistics, but never both, and given the uncertainty in our estimated threshold phylogenetic distance from the data, we disfavored such an approach.

### A Comprehensive Growth Rate Database for Amplicon Analysis

While multi-omic methods for surveying microbial communities in the environment have rapidly matured, amplicon sequencing, typically of the 16S rRNA gene, remains a cost-effective and widely used approach for assessing microbial diversity [41]. Various tools exist to link functional annotations to amplicon sequencing data, though the quality of these annotations varies widely depending on the taxonomic group and trait of interest [42-44]. Annotation quality depends on (1) the degree of trait conservation between closely related organisms, and (2) the comprehensiveness of the associated trait database used for functional annotations. Given its moderate phylogenetic conservation (see above), maximum growth rate is a suitable candidate for database-assisted functional annotation [11]. Yet, database quality remains a limiting factor.

Previously, the EGGO database of gRodon annotations addressed some of the challenges associated with functional annotation [11]. Yet, EGGO was primarily comprised of genome annotations from lab-cultivated organisms and lacked optimal growth temperature corrections, which are crucial for accurate gRodon predictions. To address these limitations, we developed an improved maximum growth rate database by 1) annotating species representative genomes from GTDB v220, which includes a majority of metagenome-assembled genomes (MAGs) and single-cell amplified genomes (SAGs) [45], 2) incorporating temperature corrections using genomic optimal growth temperature estimation software [46], and 3) applying our Phydon predictor for improved estimation. The rigorously curated GTDB provides the additional benefit of high-quality taxonomic annotations, which enhances the reliability of downstream analyses.

We ran Phydon on all 113,104 GTDB v220 species representative genomes, with 111,349 passing quality filters needed for internal gRodon prediction (*e*.*g*., having at least 10 annotated ribosomal proteins [11]). Of these, we were able to annotate 111,034 genomes with optimal growth temperature predictions from GenomeSPOT [46] and subsequently pass them through Phydon. This represents the largest temperature-corrected maximum growth rate prediction database to date. Of the 111,034 species in this database, a total of 60,869 had genomes with 16S rRNA genes present (16S rRNA recovery often fails for MAGs [47-49]), representing 17,451 genera and 191 phyla (Fig. 4). To facilitate access for researchers to this database, we provide an online tutorial alongside the Phydon package (https://github.com/xl0418/Phydon).

**Figure 4.**
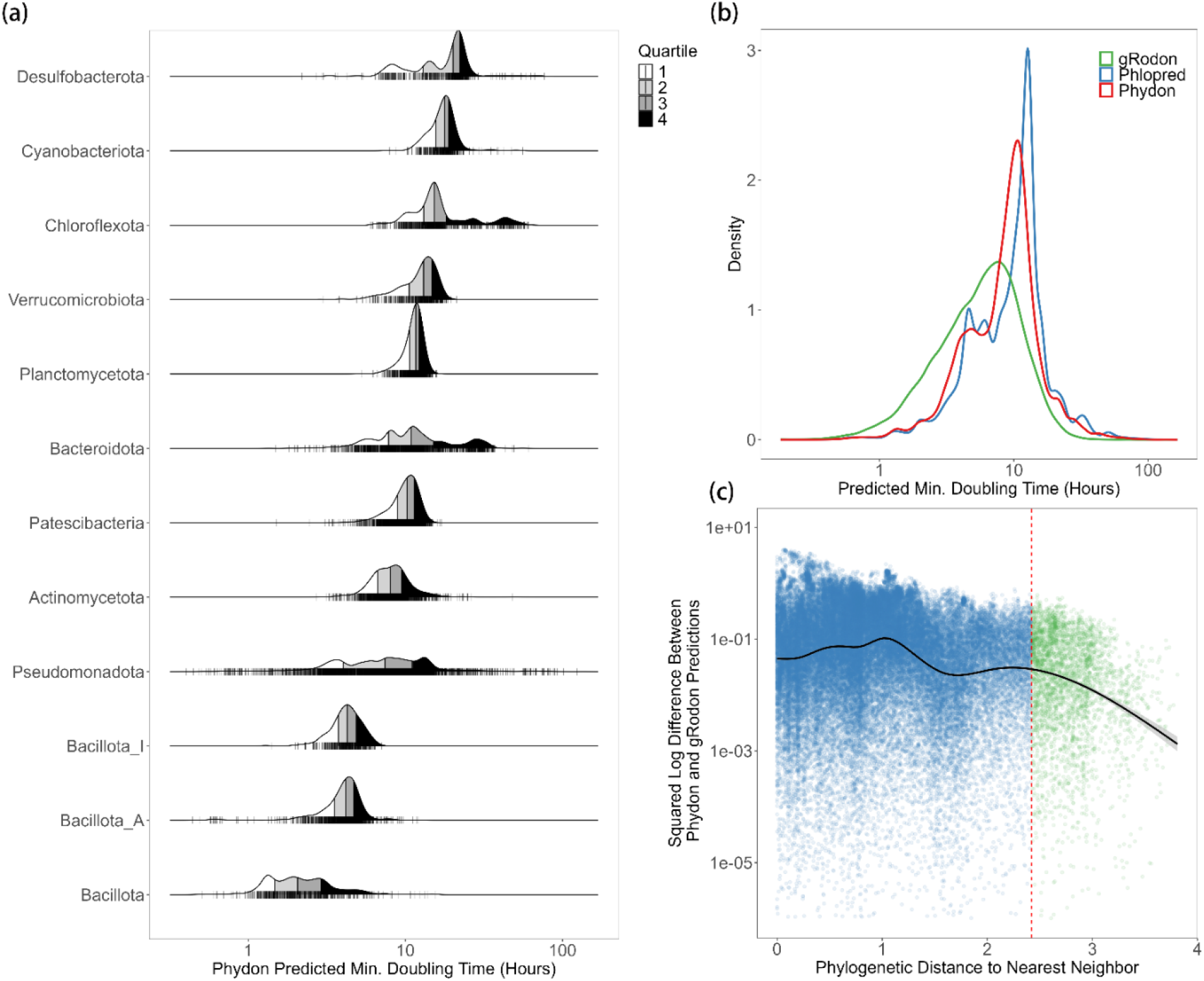
a) The distribution of the Phydon predicted maximum growth rates of major phyla (at least 1000 species representative genomes in GTDB v220; temperature corrected using genomic optimal growth temperature estimates). b) The distribution of the estimates of maximum growth rates from Phydon and gRodon and Phylopred. c) Phydon and gRodon predictions converge for distantly related organisms. Dashed vertical red line at a phylogenetic distance of 2.42 corresponding to where gRodon and phylopred have approximately equal performance.

Phydon detects major divisions in growth strategy between microbial phyla (Fig 4a), consistent with our understanding of the typical lifestyles and metabolisms associated with these groups. For example, the mostly-heterotrophic *Bacillota* (*Firmicutes*) tend to be fast-growers whereas oxygenic phototrophic *Cyanobacteria* and sulfate-reducing *Desulfobacterota* bacteria tend to be slow-growers. Our findings also reveal a clear bimodal distribution in the estimates of maximum growth rates for species in the GTDB (Fig. 4b), consistent with previous observations using the gRodon package [11]. Interestingly, Phydon is able to distinguish these major growth classes much more readily than even gRodon (Fig 4b). As anticipated, the predictions of Phydon and gRodon converge as phylogenetic distances increase (Fig. 4c), suggesting that Phydon increasingly relies on genomic factors, particularly codon usage bias (CUB), for extrapolation.

## Conclusion

Genomic and phylogeny-based methods each have distinct advantages for trait prediction under different scenarios. Specifically, the CUB-based gRodon model excels at predicting the maximum growth rates of phylogenetically distant species. In contrast, phylogenetic prediction models outperform the gRodon model for phylogenetically related species. Thus, incorporating phylogenetic information significantly enhances maximum growth rate estimation more effectively than relying solely on genomic signatures like CUB. By integrating phylogenetic context, the Phydon model achieves lower error, and more specifically lower variance than the other methods when used individually.

However, the performance of our method may be compromised when applied to taxa that have experienced rapid trait evolution. Periods of accelerated trait evolution, such as those occurring in rapidly changing environments, can weaken the predictive power of both phylogenetic models and models based on genome-scale evolutionary patterns like CUB. These models rely on the assumption of relatively stable evolutionary processes [50]. Despite this, our predictions offer a useful indication of an organism’s ecological niche, reflecting a long-term integration of evolutionary trends. It is important to interpret predicted maximum growth rates with this context in mind, as instantaneous growth rates can vary significantly over time and space depending on the local environmental conditions and the organism’s state.

A principled approach to model design involves recognizing the limitations and potential failure points of each model. By understanding where models are likely to encounter difficulties, researchers can strategically design complementary studies and integrate diverse methodologies to offset these weaknesses. Our integrated approach to trait prediction leverages phylogenetic prediction for closely related organisms and genomic model-based prediction to extrapolate to groups with limited representation in the reference database. This strategy provides a pathway for high-quality genome annotation with trait information, offering a robust solution for trait prediction for uncultured microbes.

## Methods

### The training datasets and phylogeny

We compiled a training dataset of 548 species with recorded doubling times from the Madin et al. trait database, after discarding 85 species due to unidentifiable species names in the Genome Taxonomy Database (GTDB), resulting in a total of 633 species [9] (Fig. S1). Using GTDB r220, we then extracted bacterial and archaeal phylogenetic trees [45, 51, 52]. Figure 1a presents the final phylogenetic tree for bacteria (411 species) alongside the corresponding trait distribution, while the phylogenetic tree for archaea (137 species) is presented in Figure 1b. To assess phylogenetic conservation, we calculated the pairwise differences in maximum growth rates for all species and plotted these against their phylogenetic distances. Figure S3 demonstrates that species with close phylogenetic relationships exhibit smaller trait differences, whereas more distantly related species show larger differences, highlighting the phylogenetic conservation of maximum growth rates.

Temperature is a critical factor in regulating microbial growth, and incorporating optimal growth temperature into codon usage bias (CUB)-based models can significantly enhance the accuracy of maximum growth rate predictions. We gathered data on the optimal growth temperatures of species from the Madin et al. database [9]. Of the 411 bacterial species in our dataset, 374 had recorded optimal growth temperatures, while 69 out of 137 archaeal species also had this data available. The gRodon R package features two models: one trained with temperature data and one without. To evaluate the impact of incorporating temperature on growth rate predictions, we conducted analyses using both models and compared their predictive performance to assess the enhancement provided by including optimal temperature information.

Each species in GTDB is associated with multiple genomes, with one designated as the representative genome and the remainder clustered within the GTDB phylogenetic tree. To balance information content and computational efficiency, we performed random sampling, selecting up to five genomes per species in our training set (or all available genomes if fewer than five were present) while always including the representative genome. This approach resulted in a dataset comprising 1,465 complete genomes for our 411 bacteria species and 323 genomes for 137 archaea species.

### Assessing the effect of phylogenetic relatedness on the gRodon model and phylogenetic models

Two phylogenetic prediction models were used as benchmarks to determine the conditions under which the gRodon model outperforms traditional phylogenetic prediction methods. The first phylogenetic prediction model, known as the **nearest neighbor method** (**NNM**), estimates the maximum growth rates of focal species in the testing data by averaging the maximum growth rates of the most phylogenetically related sister species in the training data. To identify the optimal number of related species for NNM, we tested groups of 1, 5, 10, 20 and 50 species. Our analysis revealed that averaging the traits of the 5 closest phylogenetic species resulted in the lowest mean squared error (MSE), although differences among group sizes were minimal (Fig. S4). Thus, we used the average trait of the 5 closest species in the NNM method.

The second phylogenetic prediction model, referred to as the **phylopred** method, predicts maximum growth rates of species using Bloomberg’s *K* statistics and Felsenstein’s independence contrast (IC) via the *phyEstimate* function from the picante package [28, 31]. This approach calculates the mean maximum growth rates of the most phylogenetically related species, with each species trait weighted according to its phylogenetic distance from the query genome. Essentially, phylopred is a weighted variant of the nearest neighbor method, where the contribution of each neighbor is adjusted based on its phylogenetic distance to the query species.

We used phylogenetically blocked cross-validation to evaluate the performance of both the gRodon model and phylogenetic prediction models [39]. We divided the phylogeny into *n* clades by cutting the tree at a uniform depth (Fig. 2). As *n* increases, the average phylogenetic distance between each pair of clades decreases (*i*.*e*., the depth at which we cut in the tree becomes shallower), meaning that for each test set the nearest clade in the training set becomes more closely related on average (Fig. 2a). We re-trained the gRodon model using the model features and structure as described in the original R code for the package on genomes from *n* − 1 clades and tested it on genomes from the remaining clade (Fig. 2bc). This process was repeated across *n* values ranging from 10 to 410 in increments of 10. When *n* = 10, there is a large phylogenetic distance between the training and test clades, while *n* = 410 represents a small minimum phylogenetic distance to the nearest clade (Fig. 2). This design created 41 test scenarios to assess model performance under varying degrees of phylogenetic relatedness.

### Phydon: Combining the gRodon model and phylogenetic prediction models

To leverage the complementary strengths of the gRodon and phylogenetic prediction models for forecasting maximum growth rates, we developed a novel combined regression model, named Phydon. This model integrates predictions from both approaches, calculating the maximum growth rate as a weighted mean of the gRodon and phylogenetic predictions. Phydon operates in two modes:

1. **Arithmetic Mean Mode**: This mode calculates the Phydon prediction as:

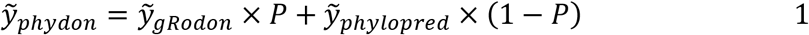

where *P* represents the probability that the gRodon model outperforms the phylogenetic models.
2. **Geometric Mean Mode**: This mode uses the geometric mean of predictions from the two models:

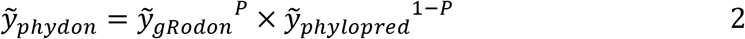

This probability *P* is determined using a regression model that considers the growth rates of the species and the phylogenetic distance (*D*) between the focal species and its nearest relatives in the training dataset. Intuitively, whether the gRodon or phylopred model should be preferred will be dependent on both how distant the query genome is from the model training data, as well as if that organism is a fast or slow grower (since model performance varies with growth rate for both approaches). The regression model, based on logistic regression using the *glm* function in R, is given by:

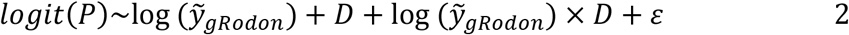

Here, *logit* (*P*) represents the log-odds of the gRodon model being superior, 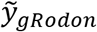 is the gRodon prediction, and *D* is the phylogenetic distance. Given that the true growth rate of the target species is unknown, the gRodon prediction serves as a proxy for growth rate. We then compared the performance of Phydon with both the gRodon model and the phylogenetic prediction models to evaluate its efficacy.

### Phydon Prediction Database

We annotated all GTDB v220 species representative genomes [45] using prokka [53] and GenomeSPOT [46]. We then passed these genomes and their optimal temperature predictions from GenomeSPOT to Phydon. For genomes with Ψ>0.6 (see [54]), indicating possible contamination, we ran Phydon using gRodon’s metagenome mode.

---

**Figure S1.**
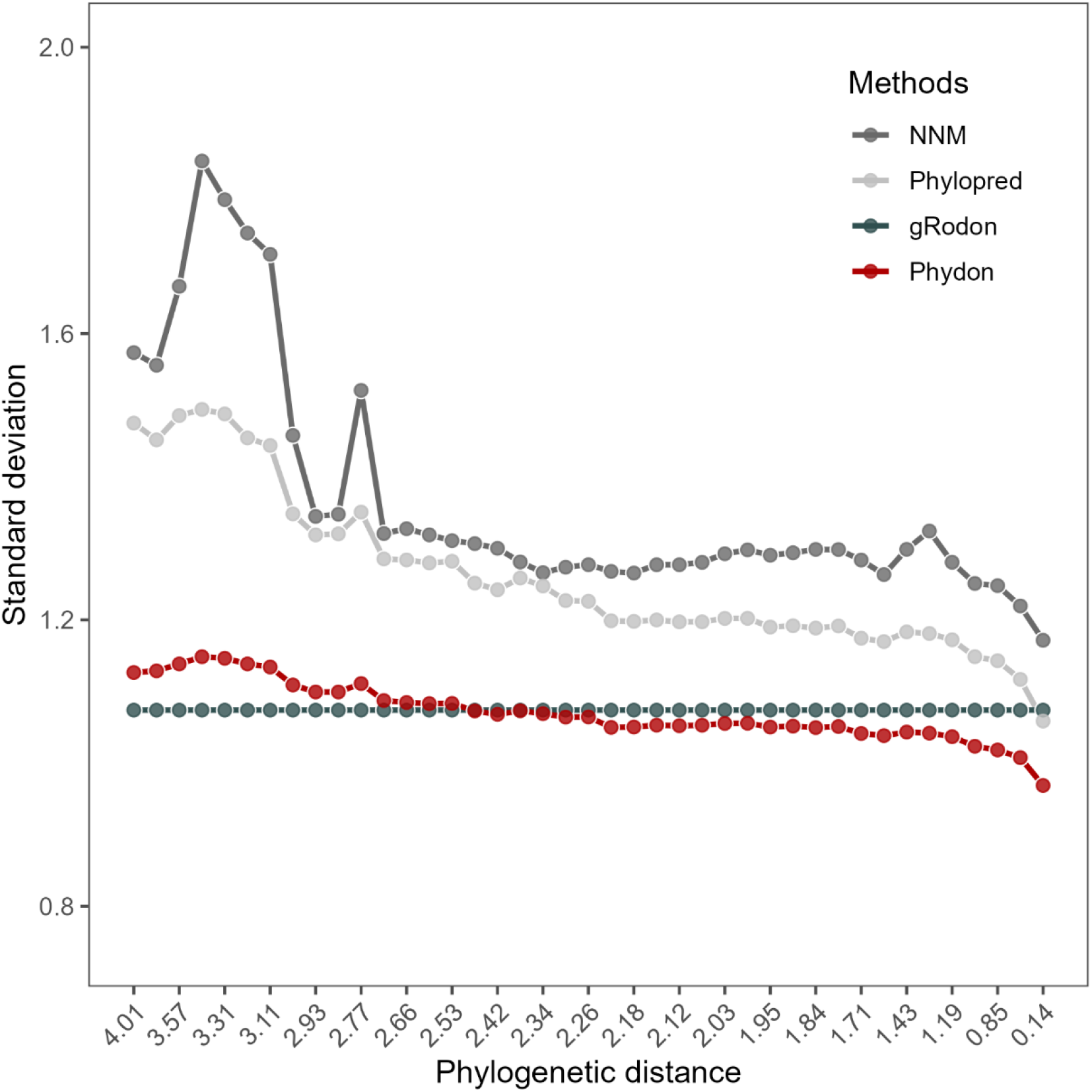
The standard deviation of the prediction error from the four prediction methods.

**Figure S2.**
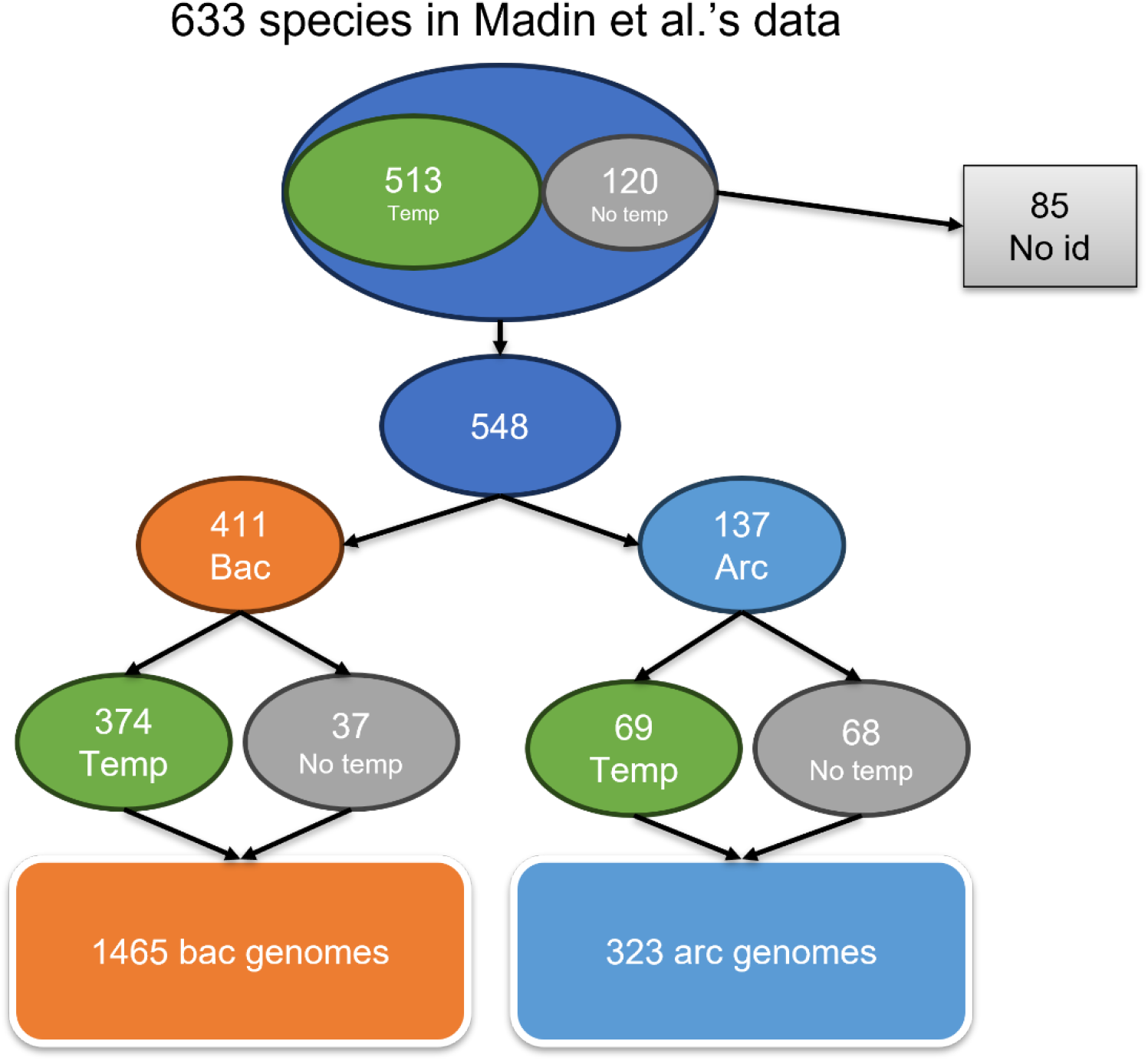
The description of Martin et al.’s data used in this study.

**Figure S3.**
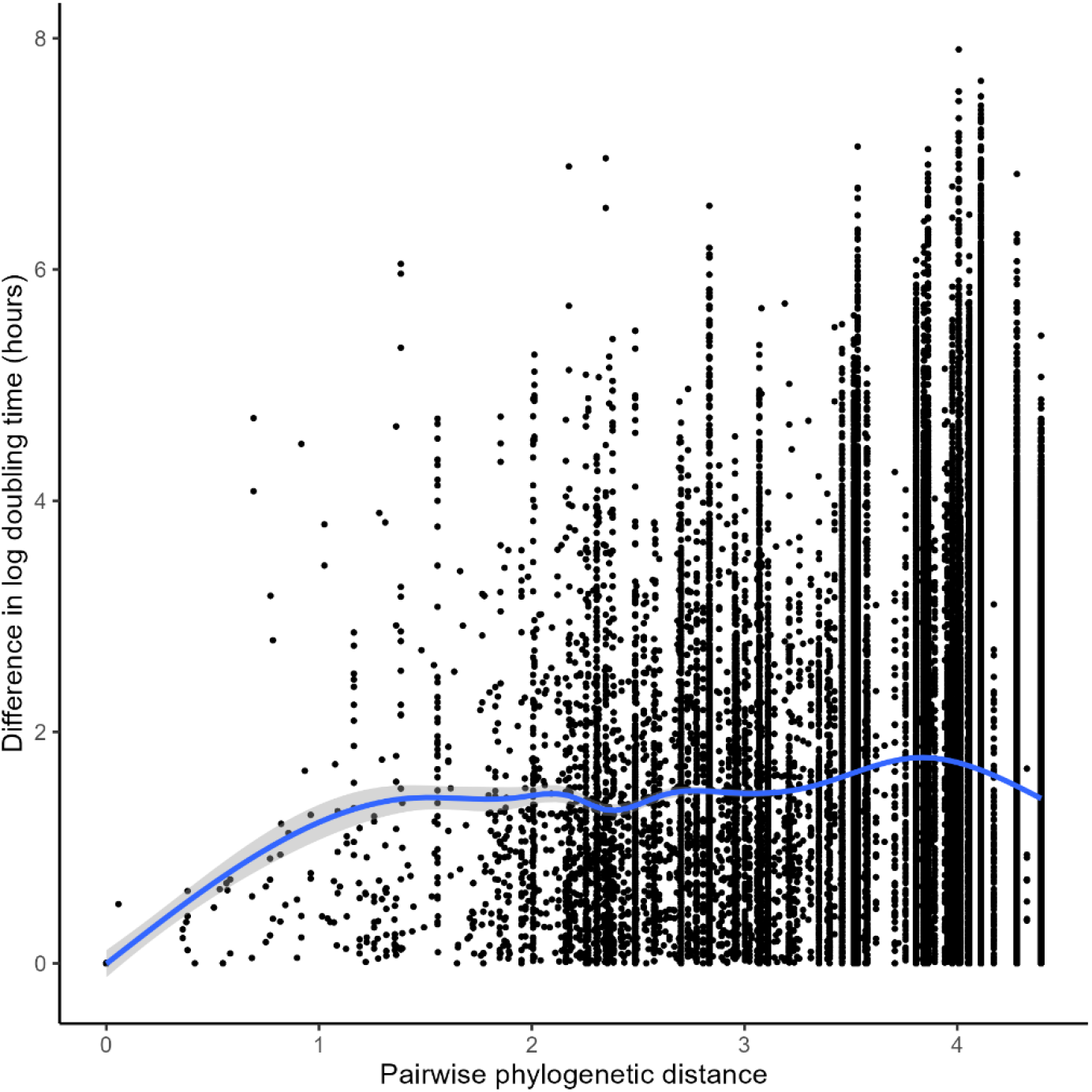
The pairwise phylogenetic distance vs. the difference in log-transformed doubling time between species.

**Figure S4.**
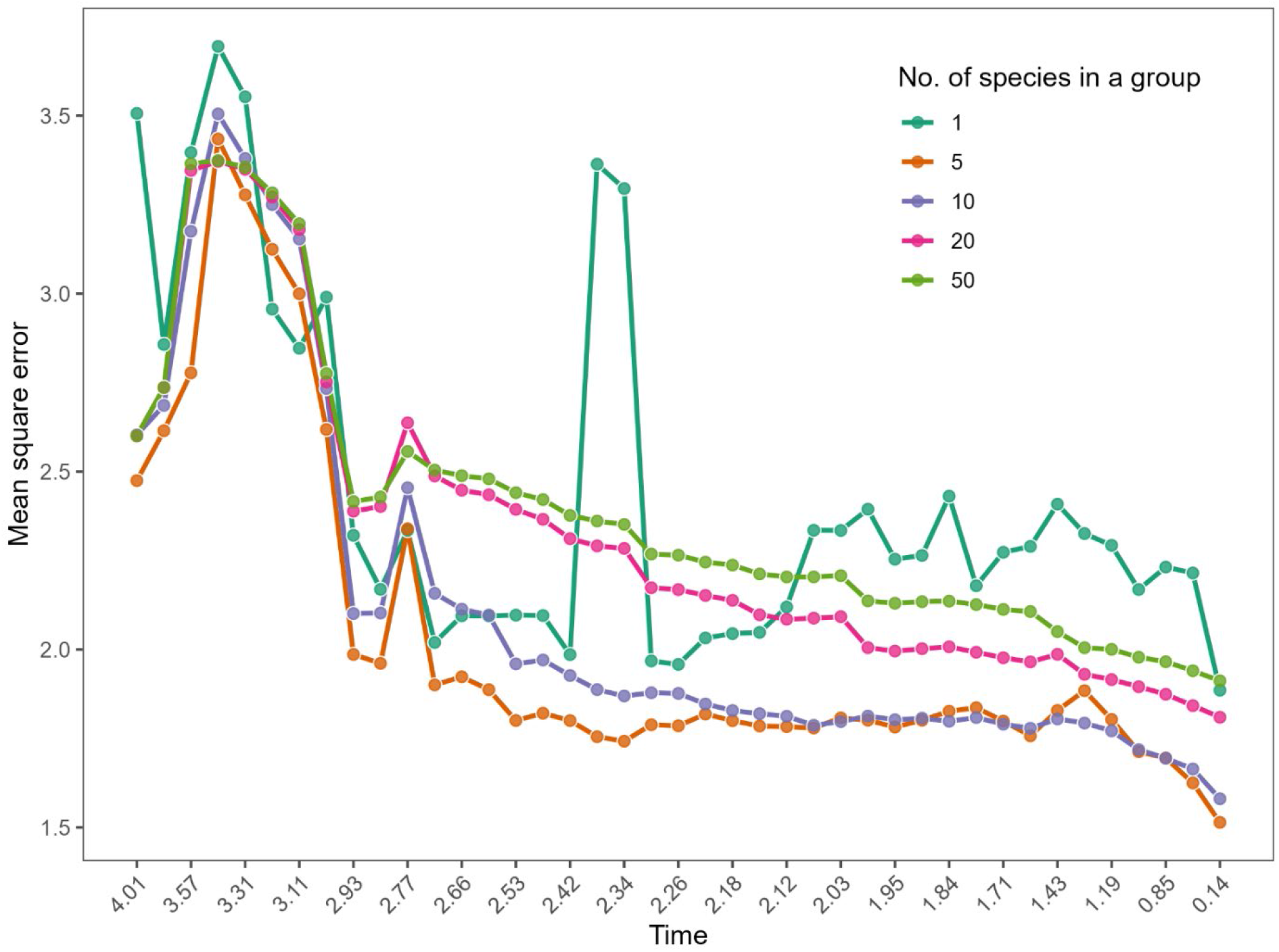
The MSE score of the NNM was calculated using different group sizes, which were used to estimate the maximum growth rates of the focal species.

